# Analysis of ovarian transcriptomes reveals thousands of novel genes in the insect vector *Rhodnius prolixus*

**DOI:** 10.1101/2020.10.22.351072

**Authors:** Vitor Lima Coelho, Tarcísio Fontenele de Brito, Ingrid Alexandre de Abreu Brito, Maira Arruda Cardoso, Mateus Antonio Berni, Helena Maria Marcolla Araujo, Michael Sammeth, Attilio Pane

## Abstract

*Rhodnius prolixus* is a Triatominae insect species and a primary vector of Chagas disease. The genome of *R. prolixus* has been recently sequenced and partially assembled, but few transcriptome analyses have been performed to date. In this study, we describe the stage-specific transcriptomes obtained from previtellogenic stages of oogenesis and from mature eggs. By analyzing ~228 million paired-end RNA-Seq reads, we significantly improved the current genome annotations for 9,206 genes. We provide extended 5’ and 3’ UTRs, complete Open Reading Frames, and alternative transcript variants. Strikingly, using a combination of genome-guided and *de novo* transcriptome assembly we found more than two thousand novel genes, thus increasing the number of genes in *R. prolixus* from 15,738 to 17,864. We used the improved transcriptome to investigate stage-specific gene expression profiles during *R. prolixus* oogenesis. Our data reveal that 11,127 genes are expressed in the early previtellogenic stage of oogenesis and their transcripts are deposited in the developing egg including key factors regulating germline development, genome integrity, and the maternal-zygotic transition. In addition, GO term analyses show that transcripts encoding components of the steroid hormone receptor pathway, cytoskeleton, and intracellular signaling are abundant in the mature eggs, where they likely control early embryonic development upon fertilization. Our results significantly improve the *R. prolixus* genome and transcriptome and provide novel insight into oogenesis and early embryogenesis in this medically relevant insect.

## Introduction

Triatomine bugs (Hemiptera, Reduviidae, Triatominae) include hematophagous species that are widespread in Latin America and have been firmly connected to the diffusion of Chagas disease. This life-threatening illness is caused by *Trypanosoma cruzi*, a protozoan parasite that is transferred from the insect vector to a human host due to the blood-feeding habit of the Triatomine species. Among these, *R. prolixus* represents a prominent arthropod vector of the trypanosome. Between 6 and 7 million people are affected by Chagas disease in Latin American countries^1^. However, several studies also reported the spread of Chagas disease in the United States of America, Canada, Europe, Australia, and Japan^2,3^. The sequencing and partial assembly of the *R. prolixus* genome were recently achieved and the latest RproC3 version is available in VectorBase^4,5^. With 702Mb of genomic DNA sequenced, it was estimated that 95% of the genome, including 15,738 genes, has been covered^4^. Despite its medical relevance, few genomic resources are available for *R. prolixus*. Over the past decade, transcriptomic analyses were carried out either using expression sequence tags or, more recently, with high-throughput sequencing techniques^6–8^. However, bioinformatic approaches coupled with molecular biology assays have recently shown that many genes are missing from the current version of the genome^9–11^. These studies highlight substantial limitations in the *R. prolixus* genome annotations, which can hinder genetic, molecular and evolutionary studies, and underscore the imperative of generating more detailed maps of the genome.

*R. prolixus* is a hemimetabolous insect that develops through five nymph stages before reaching the sexually mature adult stage^12,13^. Oogenesis in adult females is promoted by blood meals and each female can produce up to 80 eggs^14^. Females harbor two ovaries, each containing 6-8 ovarioles simultaneously^15,16^. The anterior region of the ovariole is formed by a lanceolate structure known as the tropharium that harbors the trophocytes and pro-oocytes^17^. Trophocytes are polyploid cells and are arranged in a syncytium around a central cavity known as the trophic core, while oocytes arrested in meiosis I are observed at the posterior of the tropharium. Following a blood meal and the reactivation of oogenesis, the oocytes are progressively surrounded by mitotically active follicle cells to form mature egg chambers^12,13,15,18,19^. After budding off from the tropharium, egg chambers remain connected to the tropharium through cytoplasmic bridges called trophic cords^16^. Oogenesis can be divided into three phases: pre-vitellogenesis, vitellogenesis, and choriogenesis. During previtellogenesis, RNAs, proteins, and nutrients generated by the trophocytes are collected in the trophic core and then transported via the trophic cords to the growing egg chambers^16,17,20–22^. While the growth of the oocyte is slow in the previtellogenic phase, it is rapidly promoted during vitellogenesis by fat body-derived vitellogenins and nutrients that are directly absorbed by the oocyte^23^. Once the oocyte reaches 1mm in length, the trophic cords are severed and choriogenesis begins, thus leading to the deposition of a hard chorion surrounding the oocyte. The egg displays an apparent anterior-posterior polarity with the operculum at the anterior, which is eventually displaced by the nymphs at eclosion. The dorsal-ventral axis is also readily distinguishable since the ventral region of the egg is more concave. The chorion surrounds the mature egg in *R. prolixus* and protects it against dehydration, while allowing the exchange of gases and liquids as well as fertilization^24–29^.

The genetic and molecular mechanisms that control oogenesis in the model organism *Drosophila melanogaster* have been elucidated to a great extent shedding light on key factors in important developmental processes, including germline stem cell maintenance and cystoblast differentiation, DNA damage checkpoint and DNA repair enzymes, mitosis and meiosis, and dorsal-ventral and anterior-posterior axial polarization. Furthermore, in virtually all animal species, the initial stages of embryogenesis are genetically controlled by maternally-provided molecules while the zygote is still transcriptionally inactive. Some of the mRNAs and proteins deposited by the mother in the egg can control crucial developmental decisions. In *D. melanogaster*, egg chambers of the meroistic polytrophic ovary are formed by a layer of epithelial cells surrounding the germline, which in turn, comprises 15 nurse cells and the oocyte. The nurse cells provide the *gurken*, *bicoid*, and *nanos* mRNAs, which are transported and localized into the oocyte, where they control the polarization of the dorsal-ventral and anterior-posterior axes of the egg chamber and future embryo^30^. TGF-alpha-like molecules, similar to Gurken in *D. melanogaster*, also control the establishment of the dorsal-ventral axis in wasps, beetles, and crickets^31^. In *D. melanogaster*, maternally-deposited *oskar* mRNA accumulates at the posterior of the oocyte, where it determines the nucleation of the germplasm, a specialized cytoplasm that in turn contributes to the formation of the gonads in offspring ^32^. In addition, maternal RNAs and proteins mediate the maternal-to-zygotic transition (MZT) during embryogenesis, when the developmental program switches from maternal to zygotic control ^33^. This process entails the degradation of maternal RNAs and, to a certain extent, is controlled by active mechanisms. For instance, the Smaug and Brain Tumor enzymes together with a cluster of miRNAs are responsible for maternal mRNA decay in *D. melanogaster*, while miR-430 is a key factor in maternal RNA clearance in zebrafish^34,35^. Except for a few model organisms like *D. melanogaster* and zebrafish, the genetic basis of oogenesis and early embryogenesis is still poorly understood in other animal species including *R. prolixus*.

We have recently provided an initial analysis of the transcriptome in previtellogenic stages (PVS) of *R. prolixus* oogenesis^36^. In this study, we combined the PVS datasets with newly generated transcriptomes from mature eggs resulting in more than 228 million paired-end reads across two biological replicates for each stage. Our analyses allowed us to dramatically improve the maps of genomic features in the *R. prolixus* genome and identify thousands of protein-coding transcripts and repetitive element variants. We used these data to characterize the genetic basis of *R. prolixus* oogenesis and the complement of maternally provided RNAs that drives early embryogenesis in *R. prolixus*. Finally, we have collected our results together with transcriptomic datasets from other laboratories in a mirror of the UCSC Genome Browser (Kent et al. 2002), that we named *Rhodnius* Integrated Omics Browser or RIO browser at the URL www.genome.rio/cgi-bin/hgGateway?db=rproC3.

## Results

### Improvement of the reference genome annotation with ovarian RNA-Seq datasets

We conducted a deep interrogation of the previtellogenic stage (PVS) and mature unfertilized eggs (Egg) transcriptomes, yielding in total ~228M (128M and 100M, respectively) sequence reads across two biological replicates for each stage. We previously published a partial analysis of the PVS libraries^36^, while the Egg transcriptomes were produced *de novo* for the current study. Genomic mapping of the RNA-Seq libraries was performed against the genome assembly RproC3 of *R. prolixus*^4,5^. Overall library mapping rates of the PVS and Egg conditions were on average 96% and 77.5%, respectively. The mapping rate of read-pairs aligning to multiple locations in the genome (i.e. multi-mapped read-pairs) was 12.6% on average (Supplementary Table S1). The proportion of the read-pairs unambiguously or uniquely mapped to the genome was similar in PVS and Egg samples, with 85-87% of the total mapped read-pairs. Based on these observations, the uniquely mapped read-pairs were used for a genome-guided transcriptome assembly to identify novel gene models in the genome and to improve the current gene annotations of *R. prolixus*. Preliminary quantification of the genes expressed in PVS and Egg samples was performed by assigning the uniquely mapped read-pairs to the reference gene annotation available at VectorBase (https://www.vectorbase.org/). On average, approximately 64% and 42% of uniquely mapped read-pairs of the PVS and Egg samples were respectively assigned to the reference gene annotation (column “Assigned paired-reads to reference annotation”, Supplementary Table S1). Yet, a surprisingly high percentage of uniquely mapped read-pairs, on average 36% for PVS and 58% for Egg, were not attributed to any reference genes or genomic features (Supplementary Table S1). These observations pointed to the fact that genes important for *R. prolixus* oogenesis might be missing in the current reference gene annotation. Thus, in order to expand the transcriptome landscape to include unassigned/unmapped reads, we used the uniquely mapped and unmapped reads as input for genome-guided transcriptome assembly and *de novo* transcriptome assembly approaches, respectively^37–39^. The expansion of the annotated genetic elements by the genome-guided approach provided 2,126 novel putative genes. On average, the number of genes overlapped by more than 40 read-pairs increased by 23% in PVS and Egg (Fig. 1a). It is worth noting that we adopted a conservative approach; a cutoff of 10 overlapping read-pairs is generally used to identify a new gene. For 1,659 novel genes (78%), we identified 1,204 complete ORFs, 365 (5’ or 3’) partial ORFs, and 90 internal ORFs (Supplementary Table S3). Although the remaining 476 new transcripts display small ORFs, we cannot rule out that they represent functional long non-coding RNAs (Supplementary Table S3). Interestingly, the newly annotated genes often encode putative orthologs of *D. melanogaster* proteins that are firmly linked to important developmental processes. Some notable examples in the displayed super-contigs are the *CREB-regulated transcription coactivator* (*Crtc*), the transcription factor *zelda* (*Zld*), the type II transmembrane protein *Star* (*S*), the cadherin *kugelei* (*Kug*), the homeodomain transcription factor *HTGX*, and the BMP regulator *Magu*. For easy visualization and navigation, we uploaded our datasets in a modified version of the UCSC genome browser^40^, that we named *Rhodnius* Integrated Omics or RIO browser (Fig. 1b). To facilitate the navigation, the RIO browser offers the possibility to use the name or acronym of the *D. melanogaster* ortholog to identify and visualize the features, cross-reference databases, functional classification and gene ontology of a *R. prolixus* gene.

**Figure 1.**
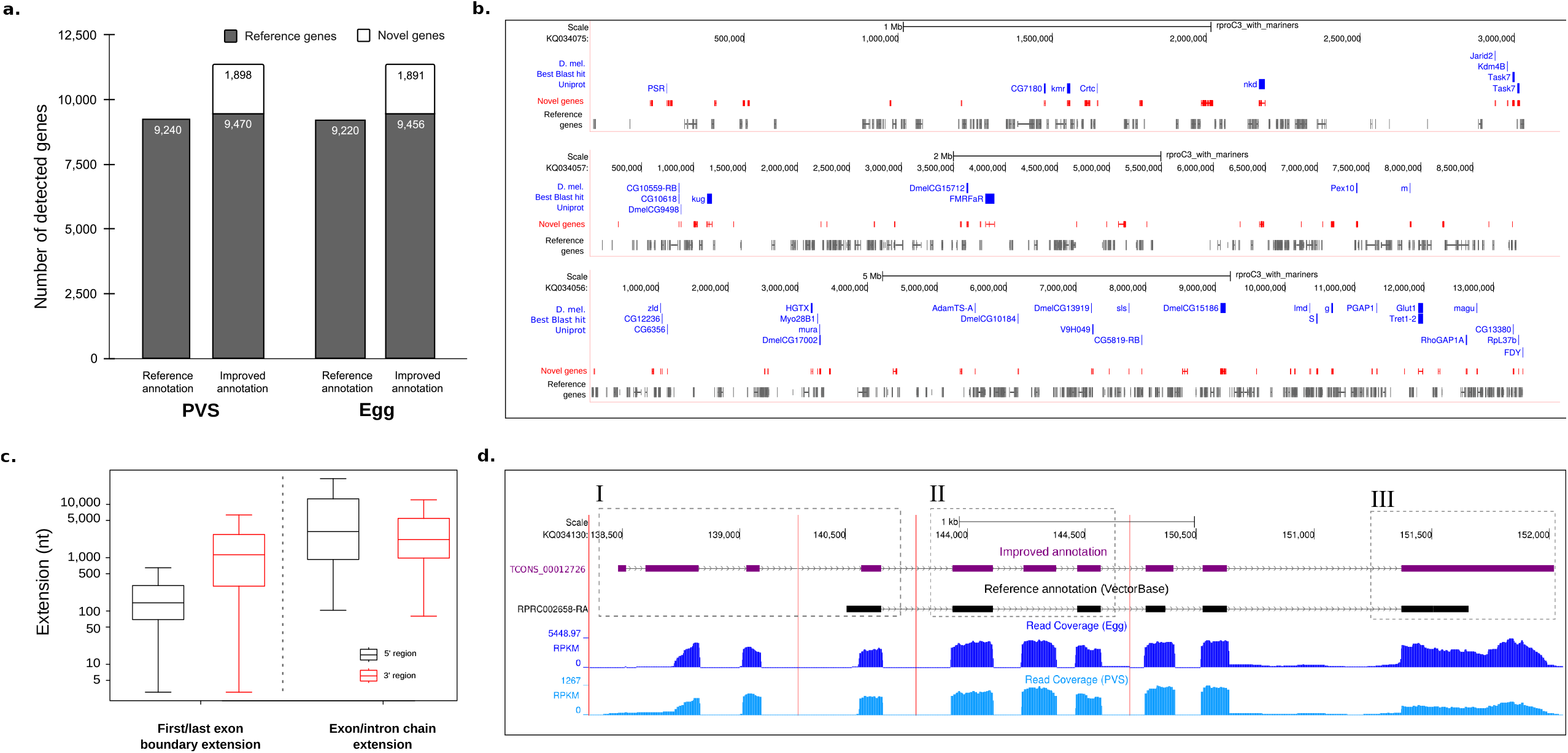
Improvement of genome annotations in *R. prolixus*. (a) Number of genes with at least 40 read-pairs overlapping the reference and improved transcriptome annotation. (b) Representative *R. prolixus* super-contigs with previously annotated genes (grey), novel genes (red), and their *D. melanogaster* orthologs (blue). (c) Distribution of the extension lengths at the 5’ (black) and 3’ (red) ends. (d) (I) Extension of the genomic coordinates of the gene RPRC002658 at the 5’ end due to the prediction of the alternative splice variant TCONS_00012726 (purple); (II) Prediction of a skipped internal exon compared to the reference transcript RPRC002658-RA (black); and (III) extension of the genomic coordinates at the 3’ end of the reference exon. Light and dark blue plots represent the read coverage of the samples PVS and Egg, respectively.

In our previous study, we noticed that the annotation of certain genes (e.g. the *R. prolixus vasa* and *PIWI* genes) was imprecise as their annotated 5’ and 3’ends did not match the gene boundaries revealed by our RNA-Seq data^36^. We, therefore, sought to provide a more detailed map of the gene structure in *R. prolixus* by genome-guided assembly of our PVS and Egg datasets. We were able to extend 1,960 genes at their 5’ ends, 1,473 genes at their 3’ ends, and 5,649 genes at both the 5’ and 3’ ends (Fig. 1c). In addition, we found approximately two new transcript variants each for 6,845 genes of the reference annotation. An assembled transcript was considered a novel isoform when it displayed (i) a multi-exon structure with at least one junction match with a reference gene in the same strand, (ii) minimum abundance of 40% of the most abundant isoform (i.e. the major isoform) of the gene and (iii) CPM > 1 at least in one condition. Figure 1c shows two types of gene extensions, one occurring at the 5’/3’ genomic coordinates of the first/last exon of an annotated gene and the other that occurs due to the prediction of new chains of exons and introns for the gene (i.e. new isoforms). The 5’ and 3’ ends were extended by a median of 569bp and 1800bp, respectively (Supplementary Table S2). By extending the annotated coding sequence (CDS), we found 78 novel complete Open Reading Frames (ORFs) in the reference transcriptome. This also improved 1,266 5’-incomplete CDSs by adding a start codon and 429 CDSs with a valid stop codon (Supplementary Table S3). A clear example of our transcriptome improvements is represented by the RPRC002658 gene encoding an uncharacterized *R. prolixus* protein (Fig. 1d). Our analyses allowed us to extend the 5’-end of this gene by approximately 2Kb (Fig. 1d, box I) due to the prediction of the novel TCONS_00012726 transcript isoform. This alternative splice variant also revealed an extra internal exon that was previously not described in the reference transcript RPRC002658-RA (Fig. 1d, box II). Finally, we were able to extend the last exon of the gene by approximately 362bp (Fig. 1d, box III). Other clear examples of the improvements that we introduced in the *R. prolixus* are represented by the genes RPRC001867 (CRAL_TRIO_N domain-containing protein, ortholog gene of *D. melanogaster* gene *cralbp*), RPRC004178 (Uncharacterized protein) and RPRC014580 (ADF-H domain-containing protein, ortholog gene of *D. melanogaster* gene GMF).

Despite the substantial expansion of the RproC3 transcriptome by genome-guided assembly, increasing the percentage of the read-pairs overlapping genomic features by up to ~40% (Table 1), 28.7% of the reads in the Egg and 4% in the PVS datasets did not map to any genomic regions (Supplementary Table S1). However, we reasoned that the stark difference between the percentage of unmapped reads in the two datasets is likely due, in great part, to a small number of genes whose RNA levels are higher in Egg versus PVS stages, given that the egg is not transcriptionally active and its RNA complement originates from the PVS. To probe this hypothesis, we applied a *de novo* assembly approach to the unmapped reads, which initially yielded 38,856 contigs for PVS and 19,390 contigs for Egg samples. After low-expression filtering, sequence deduplication, and quality contig evaluation steps (see Methods), we obtained 1,993 non-redundant protein-coding transcripts (Supplementary Table S5). However, upon closer inspection, we noticed that three contigs were accountable for >80% of the unmapped reads (Supplementary Table S4 and S5; Supplementary Fig. S1). One of these contigs encodes the *R. prolixus* Rp30 eggshell protein^41^. This abundant protein has been already biochemically characterized and a partial cDNA for this protein had already been observed in transcriptomic libraries derived from the *R. prolixus* digestive tract and ESTs datasets^6,7,41^. Our approach provides the missing sequences of the transcript and the full-length ORF. The remaining two contigs encode predicted proteins that do not share similarity with known proteins in the NCBI databases, but are particularly abundant in mature eggs. Additional studies will be required to understand whether these genes are involved in the maturation of the egg chamber, like Rp30, or they are implicated in early *R. prolixus* embryogenesis.

**Table 1.** Statistics of the read assignment to the reference and expanded transcriptomes.

|  | Genome-guided assembly |  | *De novo* assembly |  | Total assigned read pairs |  |
| --- | --- | --- | --- | --- | --- | --- |
| Samples | Uniquely mapped read pairs | Assigned to improved gene annotation | Unmapped read pairs | Assigned to *De novo* transcriptome | To the reference transcriptome | To the improved transcriptome<sup>1</sup> |
| PVS_1 | 31,651,452 | 26,759,942<br>(83.6%) | 1,565,561 | 871,556<br>(55.7%) | 20,314,277<br>(63.4%) | 27,631,498<br>(83.2%) |
| PVS_2 | 21,166,732 | 17,784,143<br>(83.4%) | 1,028,565 | 532,938<br>(51.8%) | 13,931,026<br>(65.3%) | 18,317,081<br>(82.5%) |
| Egg_1 | 17,400,997 | 15,369,913<br>(84.8%) | 6,360,014 | 5,337,815<br>(83.9%) | 8,049,949<br>(44.4%) | 20,707,728<br>(87.2%) |
| Egg_2 | 14,258,924 | 12,500,145<br>(84.8%) | 4,722,559 | 4,009,233<br>(84.9%) | 5,918,720<br>(40%) | 16,509,378<br>(87%) |

<sup>1</sup>“Expanded transcriptome” refers to the improved annotation and final putative *de novo* transcripts.

### Stage-specific gene expression during *R. prolixus* oogenesis

With our expanded version of the *R. prolixus* transcriptome, we wondered what genes are expressed during oogenesis in this species. Our quantification of the Egg datasets shows that the transcripts of 11,127 genes are present in the mature unfertilized egg (Supplementary Table S6). In *R. prolixus*, RNAs and proteins synthesized in the tropharium are transported into the oocyte via the trophic cord, while the oocyte is transcriptionally quiescent. We then performed differential expression analysis by comparing the PVS and Egg datasets in an attempt to identify stage- and tissue-specific gene expression patterns. Differentially expressed genes (DEGs) were selected according to DESeq2^42^ to identify reliable fold changes between the PVS and Egg datasets, where |fold-change| ≥ 2.5 and adjusted p-value ≤ 0.01^42^ were set as the thresholds for significantly differentially expressed genes (Fig. 2a). Here, we refer to genes as “down-regulated” or “up-regulated” if their steady-state RNA levels are respectively lower or higher in eggs compared to PVS stages (Supplementary Table S6). Differential gene expression analysis resulted in 1,480 DEGs (Fig. 2a). Of these, 851 (57.5%) genes appeared to be expressed at high levels or specifically in the PVS stage, while no expression beyond background levels was detected in Egg datasets. Surprisingly, however, 629 DEGs, which is 42.5% of the total, appeared to be highly enriched in the Egg samples. Novel putative genes represent 29% of the total DEGs, with 9.9% up-regulated and 19.1% down-regulated in PVS versus Egg stages, respectively (Fig. 2b).

**Figure 2.**
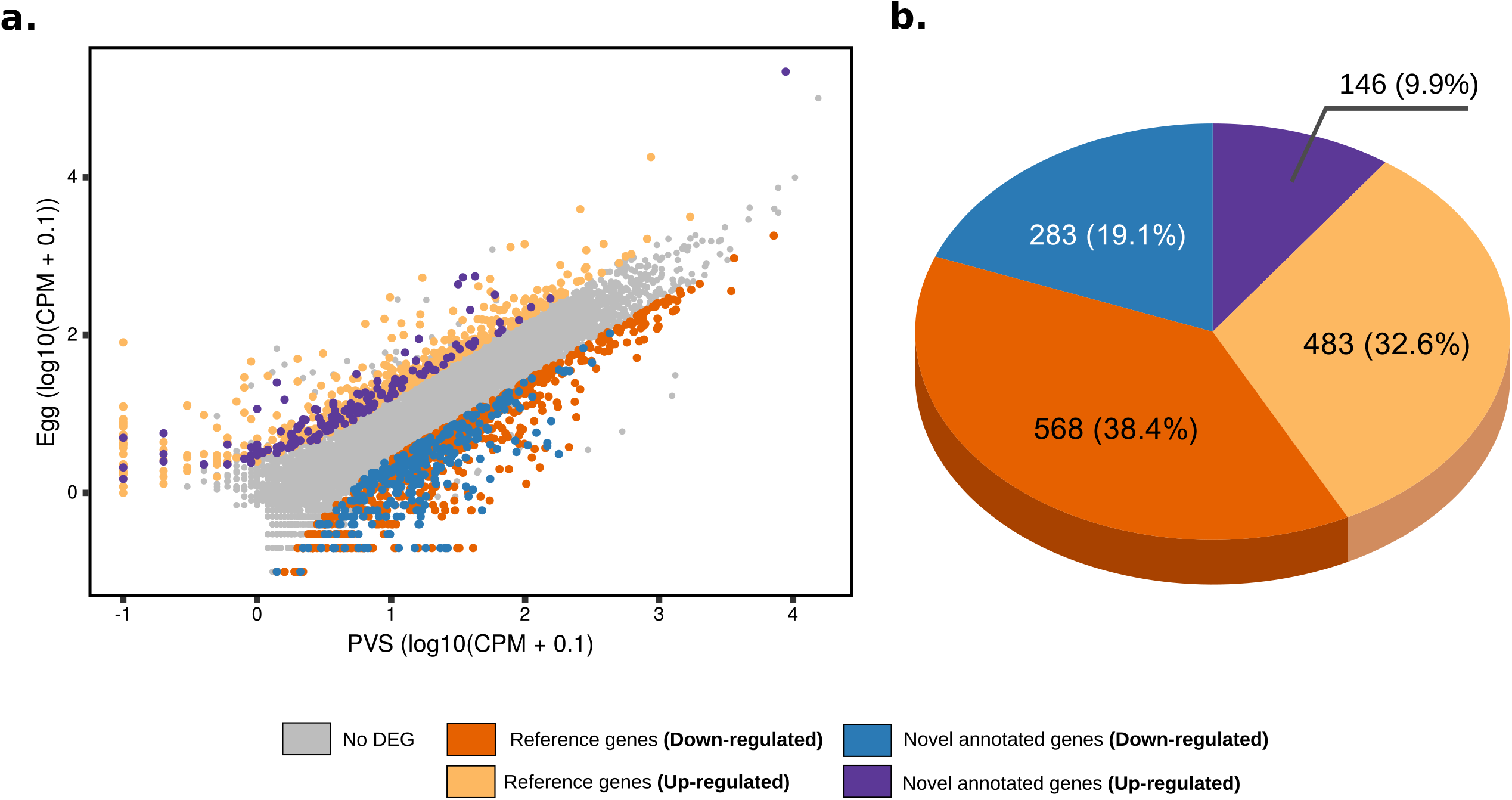
Differential expression analysis of previtellogenic stage and mature egg datasets of *R. prolixus*. (a) Scatter plot of the gene expression normalized by the average CPM (Counts Per Million, in log10). Y axis shows Egg datasets, while X displays the PVS datasets. Upregulated reference (pink) and novel (magenta) genes and downregulated reference (orange) and novel (blue) genes are displayed. Genes with no differential expression in the two stages are shown (grey) (b) Composition of the total differentially expressed (DE) genes using the improved gene annotation.

Consistent with our previous study^36^, among the top 25 genes that are highly represented in the PVS transcriptome, 15 encode ribosomal proteins (Supplementary Fig. S2). After removing these genes from the list, the most differentially expressed genes in this stage code for components of the transcription and translation machinery (Fig. 3a). Conversely, we find 13 genes in the top 25 expressed in eggs encoding uncharacterized proteins along with evolutionarily conserved proteins (Fig. 3a). On top of this list, we find the XLOC_000150 transcript that might represent a long non-coding RNA. Some examples are given by putative orthologs of the transforming growth factor-beta (TGF-β)-induced protein ig-h3 (RPRC001419), the activating transcription factor 4 (RPRC010216), a lysosomal-associated transmembrane protein (RPRC006528), and a PMSR domain-containing protein (RPRC002388), among others. Enrichment analysis of the differentially expressed genes with g:Profiler^43^ found 35 and 54 GO terms enriched (Benjamini-Hochberg FDR ≤ 0.05) in the PVS and Egg samples, respectively (Supplementary Table S7). Figure 3b shows ten major functionally grouped networks of these enriched GO terms produced by ClueGO^44,45^. This approach yielded four groups with 29 enriched terms in PVS (Supplementary Fig. S3a) and 12 groups with 54 enriched terms in Egg samples (Supplementary Fig. S3b). Our improved transcriptome confirms previous observations that the most expressed reference genes in *R. prolixus* PVS encode putative proteins with homology to *D. melanogaster* proteins related to ribosome, translation, and splicing ^36^, but it also adds enriched processes related to the P-bodies, the extracellular region, and the collagen trimer (Supplementary Table S7; Supplementary Fig. S3a). Processing bodies or P-bodies are eukaryotic cytoplasmic structures that evolutionarily conserved in organisms as distant as flies, worms and mammals^46^. In *D. melanogaster*, P-bodies are organized around the nurse cell nuclei within the egg chambers, where they form a membrane-less organelle known as “nuage”. The nuage and the P-bodies share a number of enzymatic activities and have been linked to a different molecular mechanisms including RNA decay and RNA interference. Among the genes enriched in the mature unfertilized eggs, we found protein-coding genes associated with steroid hormone receptors, cellular organization, kinase activity, transmembrane transport, protein binding, and GTPase binding. These results suggest that the mother provides a transcriptomic repertoire encoding hormone receptors (e.g. Nuclear hormone (NR) LBD domain-containing protein and ecdysone response nuclear receptor e75a), enzymes (e.g. Diacylglycerol kinases and Guanylate cyclase) and proteins with GTPase binding domains (e.g. Ran\Rab binding function and Rho GTPase exchange factor activity) critical for organismal growth, microtubule organization, cell polarity, developmental transition and regulation during the period of nutrient starvation, and transcriptional quiescence of early embryogenesis.

**Figure 3.**
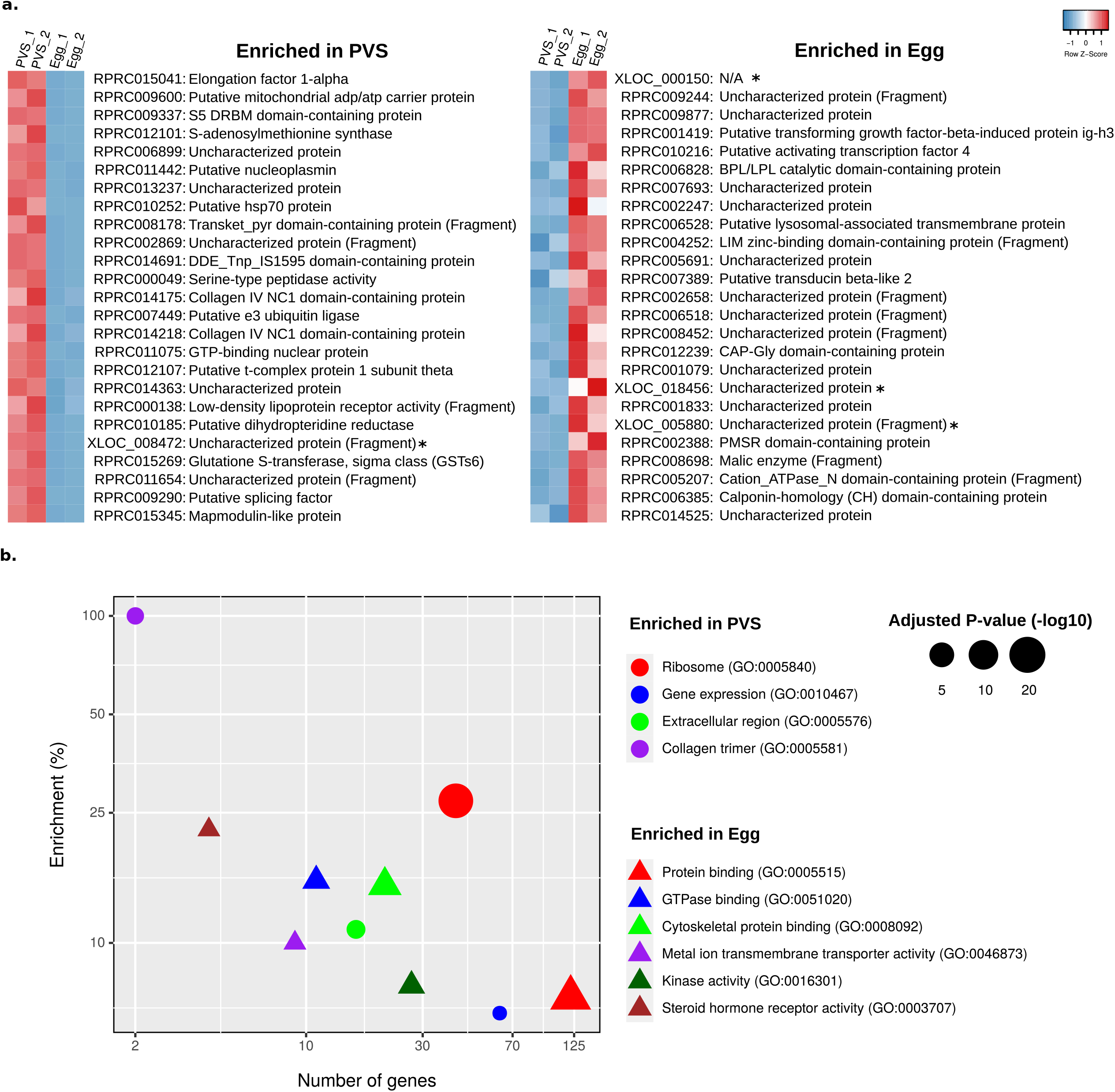
Functional categories enriched in previtellogenic stage and mature eggs. (a) Left: Top 25 differentially expressed genes (non-ribosomal-related protein) ordered by average normalized expression (CPM) for PVS, and the encoded putative protein according to the Uniprot database. Right: Top 25 most differentially expressed genes at the Egg stage. Novel genes (identifiers with suffix “XLOC” and asterisks, *) were assigned to protein products by BLAST sequence similarity. “N/A” indicates that a significant hit was not obtained (e-value ≤ 0.05). (b) Enriched GO terms representing 10 functionally related groups identified by ClueGO in the PVS (circle) and Egg (triangle) conditions. Enrichment (y-axis) is given by the number of genes in the input related to a GO term (x-axis) divided by the total number of genes associated with this GO term in %.

### Key ovarian genes of *D. melanogaster* are conserved in *R. prolixus* and expressed during oogenesis

We sought to determine the evolutionary conservation and stage-specific expression levels of the *R. prolixus* orthologs of *D. melanogaster* genes that exert critical roles in oogenesis and early embryogenesis (Fig. 4). Therefore, we focussed on six cellular and developmental processes: germline stem cells, DNA damage checkpoint and repair, mitosis/meiosis, germ granules/pole cells, axial polarization, and RNA decay ^35,47–49^. For each process, we selected well-characterized *D. melanogaster* genes, searched for the putative *R. prolixus* orthologs using BLAST tools, and measured their expression levels in PVS and Egg conditions using our RNA-Seq datasets (Fig. 4 and Supplementary Table S8). Interestingly, we find that factors involved in *D. melanogaster* germline stem cell maintenance and cystoblast differentiation are conserved in *R. prolixus* even though this cell type is not observed in the adult ovary. The expression of the *R. prolixus* orthologs of *pumilio* (*pum*) (RPRC000720), *decapentaplegic* (*dpp*) (RPRC000401), *mad* (RPRC008357), *benign gonial cell neoplasm* (*bgcn*) (RPRC008118), *medea* (RPRC003748), *glass bottom boat* (*gbb*) (RPRC013357), *zero-population-growth* (*zpg*) (RPRC014101), *hedgehog* (*hh*) (RPRC012384) and *Notch* (RPRC008058) is readily detected in both PVS and Egg datasets (Fig. 4a). The differentiation of the *D. melanogaster* germline stem cells requires the activity of *bag-of-marbles* (*bam*), a partner of BGCN in cystoblast differentiation. However, we did not find a *bam* ortholog in the *R. prolixus* genome, or in the improved transcriptome. Interestingly, *dpp* transcripts are detected only in PVS and do not accumulate in the egg.

**Figure 4.**
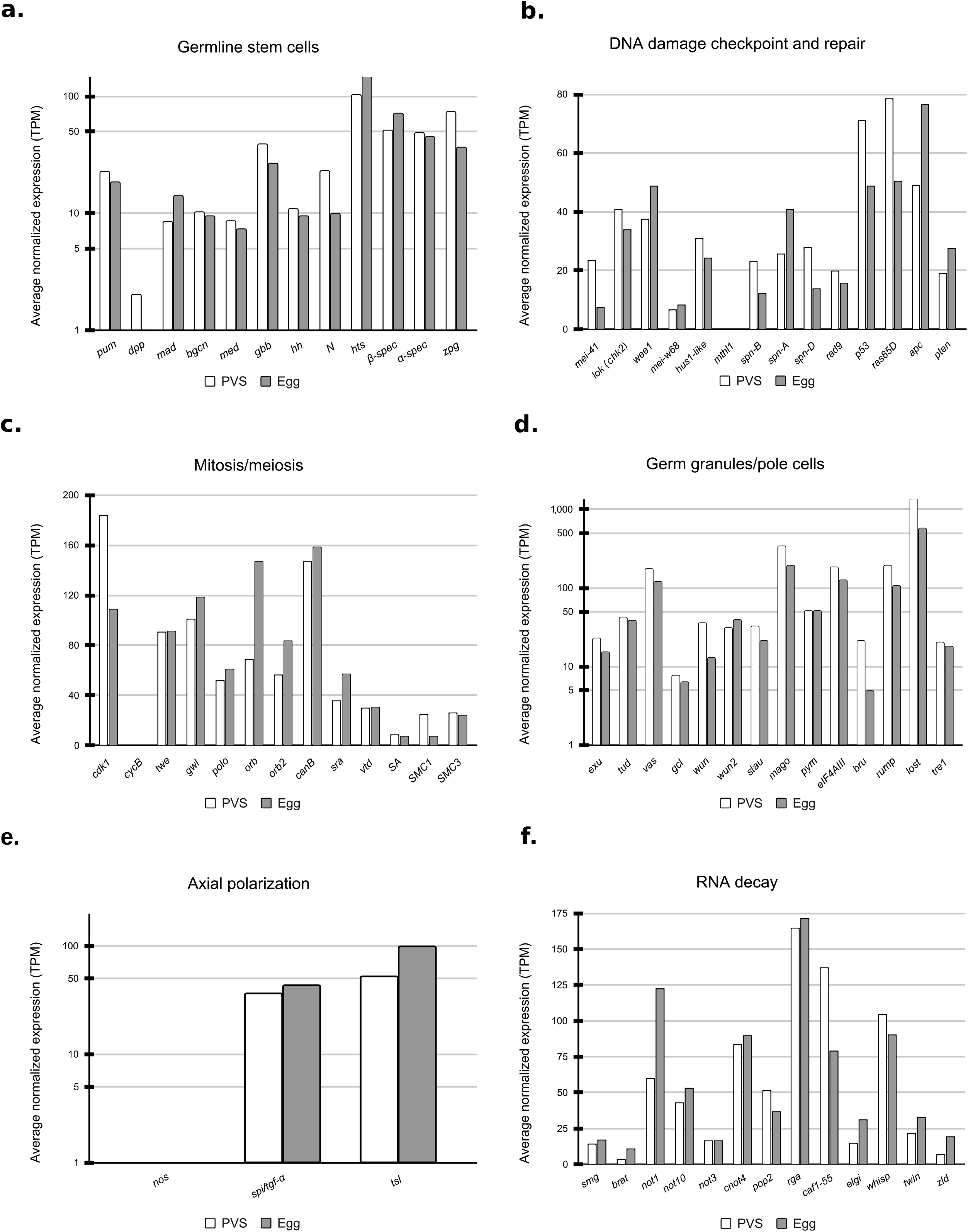
Gene expression levels of the *R. prolixus* orthologs of *D. melanogaster* genes involved in oogenesis and early embryogenesis. (a) Genes involved in maintaining germline stem cells. (b) Genes related to DNA damage checkpoint and repair processes. (c) Genes associated with mitosis and meiosis processes. (d) Essential genes for the germ granules and pole cells (e) Genes related to the axial polarization process. (f) Genes associated with RNA decay process. Y-axis displays expressions levels are normalized by Transcripts Per Million (TPM).

We also examined the expression profile of genes involved in meiosis and mitosis, including *cdk1* (RPRC009342), *cyclin B* (*cycB*) (RPRC012735), *twine* (RPRC008472), *greatwall* (RPRC005266), *polo* (RPRC002498), *orb* (RPRC006637), *orb2* (RPRC001021), *calcineurin B* (RPRC008393), *sarah* (RPRC006424), *verthandi* (RPRC005603), *stromalin* (RPRC005579), *SMC1* (RPRC009822) and *SMC3* (RPRC009208). All of the genes appear to be expressed in PVS and deposited in the mature eggs with one notable exception of *cycB*. We find two putative orthologs of *cycB* in the *R. prolixus* genome, RPRC012735 and RPRC001188, whose putative protein products share 43% and 37% amino acid sequence identity, respectively, with the *D. melanogaster cycB* gene product (Supplementary Table S8). Yet, neither is expressed at detectable levels in *R. prolixus* ovaries, nor do we find a closer ortholog in our improved transcriptome. Given that *cycB* expression is downregulated in endocycling cells like *D. melanogaster* nurse cells, and because most of the PVS tissues in *R. prolixus* are occupied by giant polyploid nuclei, it is possible that the expression of *cycB* genes in *R. prolixus* ovaries is below detectable levels.

A set of evolutionarily conserved genes is dedicated to the DNA damage checkpoint and repair in organisms ranging from yeast to mammals. DNA damage can be introduced by UV light and mutagenic agents, but DNA double-strand breaks (DSBs) introduced by *mei-W68*, the *D. melanogaster* ortholog of yeast *spo11*, are also required for meiotic recombination. DNA damage and DSBs are sensed and repaired by a set of genes including *mei-41* (RPRC004298), *chk-2* (RPRC005512), *wee1* (RPRC013702), *mei-w68* (RPRC004962), *hus1-like* (RPRC004826), *methuselah1* (*mthl1*) (RPRC001933), *spn-A* (RPRC006488), *spn-B* (RPRC010106), *spn-D* (RPRC000377), *Rad9* (RPRC001146), *p53* (RPRC003641), *ras85D* (RPRC008553), *apc* (RPRC002106), and *pten* (RPRC014866). The putative orthologs of these genes are present in the *R. prolixus* genome and the corresponding transcripts can be readily detected both in early oogenesis as well as in mature eggs (Fig. 4b).

In *D. melanogaster*, the germ plasm is a specialized cytoplasm that accumulates at the posterior of the oocyte. At the beginning of cellularization during embryogenesis, the germplasm is incorporated in the first cells appearing at the posterior of the embryo. These cells, known as pole cells, will later migrate during gastrulation and join gonadal somatic precursor cells to initiate the formation of the future gonad. Although there is no evidence of a germplasm and germ granules in *R. prolixus*, we do observe the expression of the putative orthologs of the following *D. melanogaster* genes that have been firmly connected to germplasm assembly and pole cell specification and migration: *exuperantia* (*exu*) (RPRC001415), *tudor* (RPRC012896), *valois* (RPRC005681), *vasa* (RPRC009661), *germ cell-less* (*gcl*), (RPRC010622), *wunen* (RPRC004485), *wunen2* (RPRC009835), *staufen* (RPRC010019), *mago nashi* (RPRC010985), *pym* (RPRC009783), *eIF4AIII* (RPRC002208), *bruno* (RPRC007498), *rump* (RPRC014343), *lost* (RPRC014809), and *tre1* (RPRC011268). All of the corresponding *R. prolixus* genes seem to be expressed early in oogenesis and accumulate in the mature eggs. The steady-state RNA levels of the putative *lost* ortholog in *R. prolixus* (>1000 TPM) suggest that this gene exerts a prominent role during *R. prolixus* oogenesis. As expected, we did not find a *R. prolixus* ortholog of the *oskar* gene, that is known to be restricted to Diptera, while, consistent with our previous findings, the *R. prolixus vasa* gene is expressed in PVS and the transcripts are deposited in mature chorionated eggs ^36^.

Axial polarization in *D. melanogaster* is dictated by maternal transcripts encoding the Bicoid and Gurken proteins, whose localized translation sets the anterior-posterior and dorsal-ventral axes, respectively, of the egg and future embryo. Neither of these two genes appears to be conserved in *R. prolixus*, although we found a *spitz/tgf-a* gene (RPRC009386) that is expressed in both the PVS and Egg datasets. Members of the TGF-alpha family were shown to control the formation of the dorsal-ventral axis in crickets, wasps, and beetles^50^. We also find putative orthologs of *nanos* (RPRC002927) and *torso-like* (RPRC006513), but not of *torso*, which are involved in the specification of the anterior-posterior axis in fruit flies, aphids and honeybees^51^.

Finally, we investigate the expression profiles of genes associated with the RNA decay pathway. The early phases of embryonic development in *D. melanogaster* and other organisms are driven by maternally contributed RNAs and proteins. However, the maternal program must be erased at the onset of MZT so that the embryo can follow a zygotically-driven transcription program. The degradation of maternal RNAs in *D. melanogaster* is mostly guided by the protein products of a group of genes comprising *smaug* (RPRC007649), *brain tumor* (RPRC007254), *not1* (RPRC005691), *not3* (RPRC003870), *not10* (RPRC001957), *cnot4* (RPRC001525), *pop2* (RPRC004491), *regena* (RPRC003111), *caf1* (RPRC015316), *early girl* (RPRC004083), *wispy* (RPRC007755), *twin/CCR4* (RPRC006991), and *string* (RPRC008472). In addition, *zelda* (RPRC000020) encodes a transcription factor that is pivotal for the onset of zygotic transcription ^52^. We find that all of these genes are expressed in PVS and their transcripts are detected in mature eggs. Unexpectedly, the levels of *smaug*, *wispy*, and *zelda* are low and might suggest that other mechanisms are in place during early *R. prolixus* embryogenesis to degrade maternal RNAs and prompt zygotic transcription. We did not find *R. prolixus* orthologs for the following key *D. melanogaster* genes in the genome or in our updated transcriptome: *fs(1)Yb*, *swallow*, *trunk*, *barentsz*, *atr/tefu*, *cup*, and *matrimony*. Despite the extensive conservation of crucial genes between *D. melanogaster* and *R. prolixus*, we also detect important differences, which might explain, at least in part, the striking differences between oogenesis and embryogenesis in these two insect species.

### Expression levels of transposable and repetitive sequences during *R. prolixus* oogenesis

Transposable elements (TE) are mobile sequences that often constitute a significant proportion of animal and plant genomes. While the biological function of these selfish sequences is still debated, it is well established that transposon mobilization is often connected to DNA damage, insertional mutagenesis, and genome instability. RepeatMasker annotation of RproC3, available at VectorBase, shows that more than half of *R. prolixus* mobilome is dominated by repetitive elements (longer than 0.5Kb) of the type Tc1-*mariner* (28%), *Helitron* (14%), *Tc1* (10%) and a class of repetitive elements labeled as *Unknown* (9.8%) (Supplementary Table S9). The prevalence of Tc1-*mariner* elements has been reinforced using *de novo* and library-based approaches for transposon detection, revealing that approximately 75% of the transposons dispersed in the *R. prolixus* genome belong to the Tc1-*Mariner* family^53^. Given the incomplete assembly of the genome and the challenges imposed by the repetitive nature of these elements, it is likely that the mobilome of *R. prolixus* is not completely characterized. Furthermore, the expression levels of these sequences have not been investigated yet in *R. prolixus*. We therefore employed the available data together with our stage-specific RNA-Seq datasets to determine the expression levels of all known transposon families in *R. prolixus* oogenesis. As for the protein-coding genes, we also compared the early (PVS) and late (Egg) stages of oogenesis to identify transposons potentially displaying differential activity in these two stages (Fig. 5). First, we calculated the RNA-Seq data coverage of the transposable elements by assigning all of the mapped read-pairs to the RproC3 RepeatMasker annotation and our improved gene annotation. Read-pairs assigned to more than one feature or genomic element class (e.g., protein-coding gene, transposable element, simple repeats) were considered ambiguous and, therefore, not included in the pie charts (less than 7% of the total assigned read-pairs) (Fig. 5a). In agreement with their relative abundance in the genome, the Tc1-*mariner* family of transposons appeared to have the highest expression levels both in PVS and in mature eggs (Fig. 5b). High expression levels are also observed for a class of “unknown” transposons and the *Helitron* family. When comparing the steady-state levels in PVS versus Egg datasets, we observed that some classes of DNA transposons, LTR and Non-LTR retrotransposons, have greater coverage in the previtellogenic stage (like the “unknown” TE class and *gypsy*), while others have greater coverage in the mature unfertilized egg (like Tc1-*mariner* and *helitron*). Despite these differences, however, it seems that all of the transposon transcripts that are generated in the tropharium are efficiently transported to the growing oocyte. We were also interested in identifying novel repetitive element (RE) variants, more specifically transposons. To do so, we performed homology searches based on hidden Markov model (HMM) profiles of transposable elements^54,55^ against our *de novo* assembled contigs. In total, homology searches resulted in 167 contigs with significant hits of non-rRNA repetitive elements. To reduce the sequence redundancy between stages, we aligned the new repetitive elements with each other, resulting in 20 non-redundant putative REs (Supplementary Table S10). We also aimed to identify whether our transposons were already characterized in the library of RproC3 repetitive elements available at VectorBase or found in published Tc1-*mariner* consensus sequences^53^. Interestingly, we found 12 non-redundant transposon consensus sequences with no significant similarity or with low alignment coverage with RproC3 repetitive elements^4,5^ or Tc1-*mariner* consensus sequences^53^ (Table 2). Abundances of these sequences are represented by the number of estimated reads normalized by the total of estimated reads (estimated CPM) for each replicate (“Rep.”) of the PVS and Egg *de novo* contigs. Our study clearly shows that a variety of transposons belonging to known or uncharacterized families are expressed in early oogenesis and their transcripts accumulate in the mature eggs.

**Table 2.**
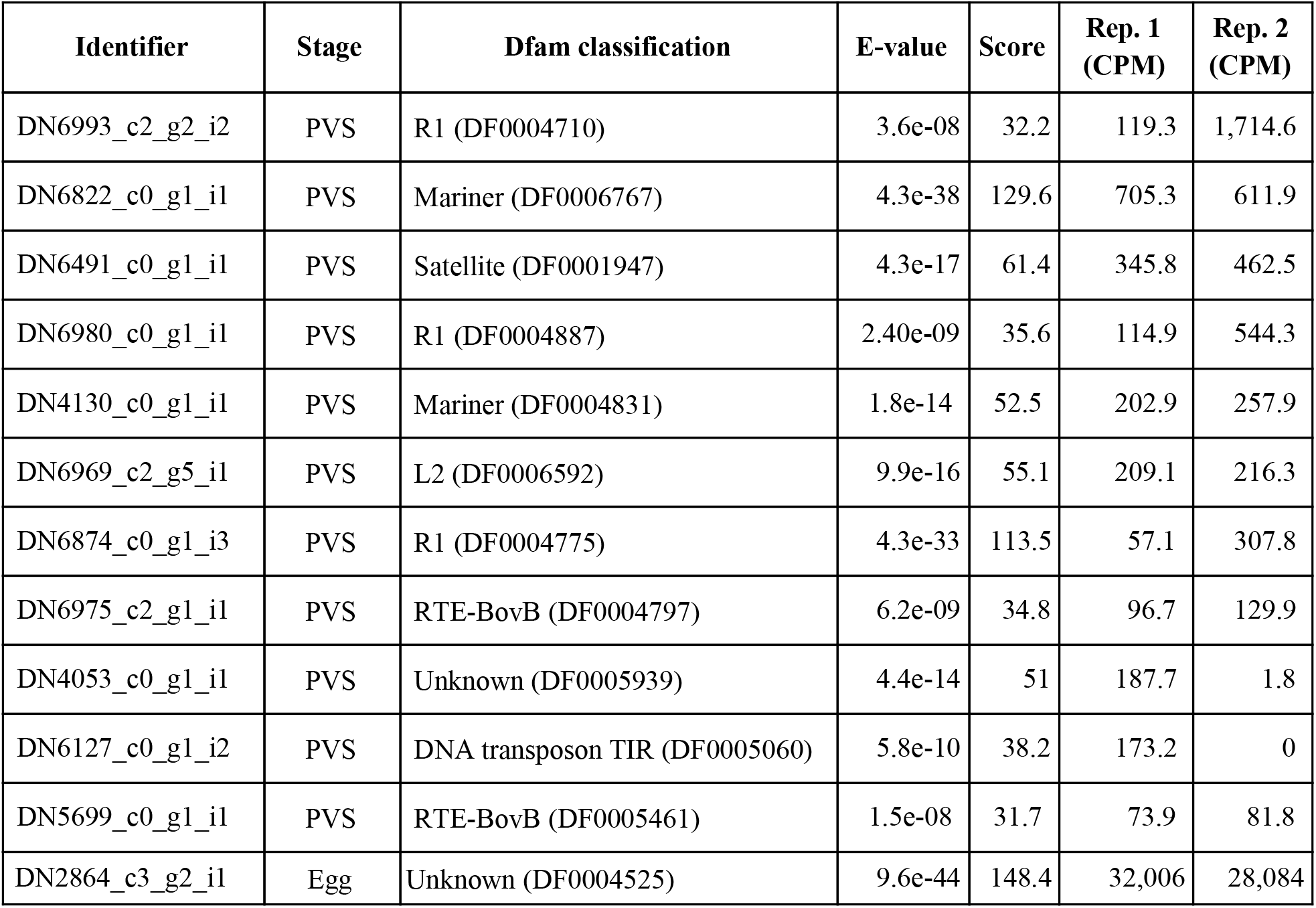
Putative repetitive elements and transposons non-redundant sequence variants.

| Identifier | Stage | Dfam classification | E-value | Score | Rep. 1 (CPM) | Rep. 2 (CPM) |
| --- | --- | --- | --- | --- | --- | --- |
| DN6993_c2_g2_i2 | PVS | R1 (DF0004710) | 3.6e-08 | 32.2 | 119.3 | 1,714.6 |
| DN6822_c0_g1_i1 | PVS | Mariner (DF0006767) | 4.3e-38 | 129.6 | 705.3 | 611.9 |
| DN6491_c0_g1_i1 | PVS | Satellite (DF0001947) | 4.3e-17 | 61.4 | 345.8 | 462.5 |
| DN6980_c0_g1_i1 | PVS | R1 (DF0004887) | 2.40e-09 | 35.6 | 114.9 | 544.3 |
| DN4130_c0_g1_i1 | PVS | Mariner (DF0004831) | 1.8e-14 | 52.5 | 202.9 | 257.9 |
| DN6969_c2_g5_i1 | PVS | L2 (DF0006592) | 9.9e-16 | 55.1 | 209.1 | 216.3 |
| DN6874_c0_g1_i3 | PVS | R1 (DF0004775) | 4.3e-33 | 113.5 | 57.1 | 307.8 |
| DN6975_c2_g1_i1 | PVS | RTE-BovB (DF0004797) | 6.2e-09 | 34.8 | 96.7 | 129.9 |
| DN4053_c0_g1_i1 | PVS | Unknown (DF0005939) | 4.4e-14 | 51 | 187.7 | 1.8 |
| DN6127_c0_g1_i2 | PVS | DNA transposon TIR (DF0005060) | 5.8e-10 | 38.2 | 173.2 | 0 |
| DN5699_c0_g1_i1 | PVS | RTE-BovB (DF0005461) | 1.5e-08 | 31.7 | 73.9 | 81.8 |
| DN2864_c3_g2_i1 | Egg | Unknown (DF0004525) | 9.6e-44 | 148.4 | 32,006 | 28,084 |

**Figure 5.**
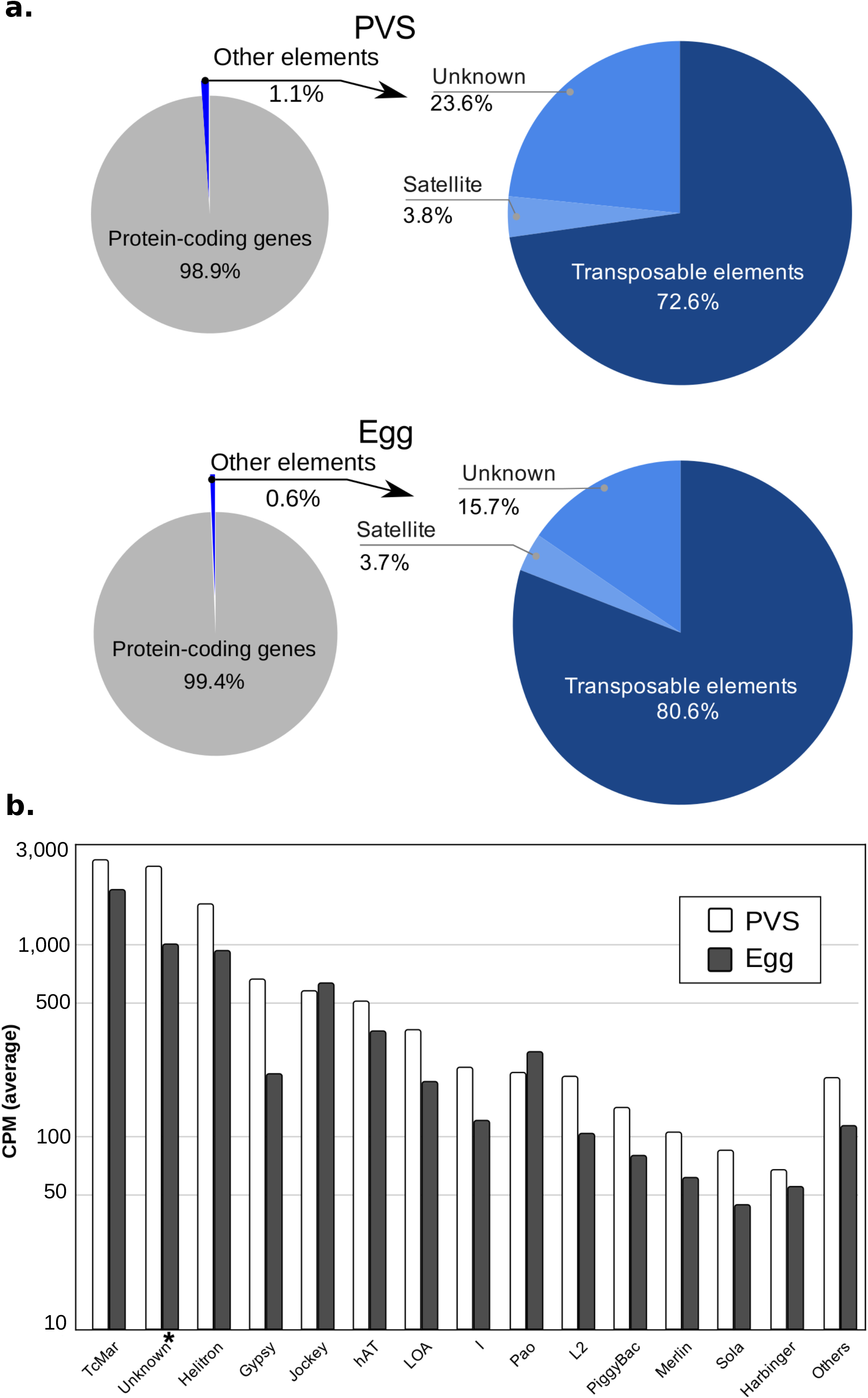
RNA levels of transposable and repetitive elements. (a) Left: pie chart shows the assignment of the multi- and uniquely-mapped paired-end reads to the protein-coding gene annotation (gray) and repetitive element regions (blue). Right: pie chart shows a more detailed distribution of mappings for satellite simple repeats, transposable elements, and unknown classes of repetitive elements. (b) Cumulative quantification of expression for each class of transposable elements longer than 0.5 Kb (kilobases) for the conditions PVS (white) and Egg (gray).

## Discussion

*R. prolixus* is a primary vector of *Trypanosoma cruzi*, the etiologic agent of Chagas disease. Sequencing of the *R. prolixus* genome in recent years provided an important resource to understand the biology of this insect. In this study, we sought to improve genome annotations and use the improved transcriptome to gain insight into *R. prolixus* oogenesis and the maternal contribution to early embryonic development. Using a combination of genome-guided and *de novo* transcriptome assembly, we were able to increase the number of putative protein-coding genes in *R. prolixus* from the 15,738 that are currently annotated in the genome (RproC3 version) to 17,864. Many of these genes likely code for critical factors in *R. prolixus* development as they include histone modification enzymes and chromatin remodeling factors, homeotic genes, signaling molecules and kinases, and metabolic enzymes, many of which are evolutionarily conserved. Furthermore, we significantly improved the 5’ and 3’ boundaries of previously annotated genes, thus shedding light on their 5’ and 3’ UTRs. Importantly, for 2,709 of these genes, we revealed the complete ORF and the encoded full-length putative protein. Finally, the paired-end nature of our libraries allowed us to determine transcript variants generated by alternative splicing events for 6,845 genes.

Based on the updated transcriptome, we investigated the complement of genes expressed in early stages of *R. prolixus* oogenesis and drive egg chamber formation and oocyte maturation, as well as the atlas of transcripts that are maternally deposited in mature unfertilized eggs. The latter genes are essential during the first stages of embryogenesis, when zygotic transcription is not yet active. The morphology and architecture of the ovarioles differ profoundly in *R. prolixus* and *D. melanogaster*. While in *Drosophila* the oocytes are produced from germline stem cells located at the anterior tip of the ovariole, this cell type seems to be restricted to nymph stages in *R. prolixus*. Instead, the anterior region of the ovariole harbors actively dividing trophocytes in this species^15–17^. Despite this difference, we find orthologs of *Drosophila* genes involved in germline stem cell specification and maintenance, like *hts*, *α-spec, bgcn* and *dpp*, in early stages of *R. prolixus* oogenesis. Particularly surprising is the expression of *hts* and *a-spec*, given that the products of these genes in *D. melanogaster* mark the spectrosome/fusome, a cytoplasmic structure that initially labels the germline stem cells, but once the cystoblast differentiates and start diving, it forms a bridge connecting the germ cells within the mitotic cyst. The spectrosome/fusome exerts an important role in oocyte specification in the fruit fly. Thus, it will be of great interest to characterize its localization and function during *R. prolixus* oogenesis, where the pro-oocytes are already specified in earlier nymph stages and germline stem cells are not detected in the adult ovary. Our analysis also shows that many *D. melanogaster* genes controlling critical processes like meiosis/mitosis, DNA damage checkpoint and repair, germ plasm assembly and pole cell formation, maintenance and migration and RNA decay are conserved in *R. prolixus*. Remarkably, the *R. prolixus* orthologs of several genes, whose expression in *D. melanogaster* is restricted to early oogenesis like *hts*, *α-spec*, *bgcn*, and *mei-W68*, are instead expressed in PVS and their transcripts deposited in mature eggs. This might represent a consequence of the particular structure of *R. prolixus* egg chambers and ovarioles. The egg chamber is exclusively formed by somatic follicle cells surrounding the oocyte and is constantly communicating with the tropharium through the trophic cords until choriogenesis begins. Differently, the egg chamber in the fruit fly is formed by a layer of follicle cells surrounding the germline, where the germline comprises the oocyte and fifteen nurse cells. Once the egg chamber buds of the germarium in *D. melanogaster*, it remains physically detached from the germarium, and the supply of RNAs and proteins for the growth and patterning of the oocyte is solely provided by the adjacent nurse cells through ring canals. Therefore, RNAs and factors expressed in early stages of oogenesis (i.e. tropharium/germarium) can be found in *R. prolixus*, but not in *D. melanogaster* mature eggs. The presence of transcripts encoding components of the RNA decay pathway, including *smaug*, *wispy*, and the CCR4/Not complex, suggest that maternal RNAs are actively degraded in *R. prolixus* to promote the maternal-to-zygotic transition. As expected, some patterning genes like *gurken* and *bicoid*, which are responsible for axial polarization in *D. melanogaster* are not detected in the *R. prolixus* genome given that they are restricted to Diptera. However, we identified a putative ortholog of *D. melanogaster spitz*, which encodes a TGF-α protein, expressed in *R. prolixus* oogenesis. Members of this family, which include Gurken, have been connected to the establishment of the dorsal-ventral polarity in crickets, wasps, and beetles^31^. Also, we did not find an obvious ortholog of the germ plasm factor *oskar*, although a *oskar* gene was identified in cricket, a hemimetabolous insect like *R. prolixus^56^*. Another patterning gene expressed in *R. prolixus* oogenesis is *nanos*. Nanos is responsible for the specification of the posterior segments in a range of distantly related insect species^50,57,58^. Although these observations suggest that the mechanisms controlling axial polarization and germplasm formation are partially conserved in *R. prolixus*, functional studies are required to gain further insights into these processes.

Our data show that the transcripts encoding a variety of ribosomal proteins rank among the top 25 genes that are expressed early during oogenesis, but are under-represented in mature unfertilized eggs. It is possible that in *R. prolixus* ribosomal proteins rather than the corresponding transcripts are deposited in the egg and used in early stages of embryogenesis. This strategy might guarantee that ribosomal proteins are readily available to rapidly assemble ribosomes. Surprisingly, we found 629 genes to be highly represented in the Egg samples, but not in the PVS. It seems reasonable to assume that these genes are activated in the tropharium in response to the blood meal, but their expression is not sustained over time. The transport of the corresponding transcripts to developing oocytes would, therefore, progressively deplete the tropharium of these RNAs while they accumulate in the egg. Among the top 20 of DEGs with high expression in mature eggs, we find a putative ortholog of *Yellow-g2* (RPRC009244). This gene has been associated with the resistance of mosquito eggs to desiccation as well as to the production of cuticle pigments in *R. prolixus^59^*. At least two genes might be activated in oogenesis to control the oxidative stress caused by the blood feeding habits of *R. prolixus*. In fact, RPRC010216 encodes a putative ortholog of the cyclic AMP-dependent transcription factor ATF-4, a key regulator in metabolic and redox processes as well as a crucial factor in the integrated stress response, and RPRC002388 encoding a predicted PMSR-containing protein. Peptide Methionine Sulfoxide Reductase (PMSR) was associated with the repair of proteins and peptides that have been damaged by oxidizing environments^60^. It is well-established that the blood-feeding habit of *R. prolixus* generates oxidative species. Hence, RPRC002388 might exert a crucial role in the repair of proteins during *R. prolixus* oogenesis and early embryonic development. Another gene that might be linked to the heme metabolism is RPRC001419 encoding a putative ortholog of the venom periostin-like protein 1 from *Pristhesancus plagipennis*, also an assassin bug like *R. prolixus*, but preying on other insects and not on mammals. *Pristhesancus* injects the venom into the prey to paralyse it and liquify the tissues to facilitate the feeding. Finally, some promising genes need to be validated by other molecular biology techniques. For instance, RRPC009877 coding for a putative ortholog of a proline-proline-glutamic acid (PPE) protein from *Mycobacterium.* The PPE protein family has been connected to cell wall remodeling and virulence in bacteria. It is noteworthy however, that the neighboring gene in the contig, namely RPRC009884, also encodes for a putative bacterial protein. It will be important to determine whether these genes are truly present in the *R. prolixus* genome or they are bacterial genes accidentally assembled in the insect genome.

Our data also reveal that a variety of transposable elements are expressed in the early stages of oogenesis and their transcripts accumulate in the developing oocyte. It was previously shown that approximately 6% of the *R. prolixus* genome is made up of transposable sequences, with the Tc1-*mariner* elements comprising 75% of the mobilome. In accordance, we find that the Tc1-*mariner* transposons display the highest steady-state RNA levels in both PVS and Egg samples compared to other transposon families. However, the *helitron* elements also appear to display comparable expression levels to Tc1-*mariner* together with a class of “unknown” repetitive sequences, even though they are less represented in the genome. Overall, transposable and repetitive sequences account for only 1.1% and 0.6% of the unambiguous reads in PVS and Egg stages, respectively. We previously showed that key components of the piRNA pathway are conserved in *R. prolixus* and are important for female adult fertility ^36^. Together with the observation that *R. prolixus* ovaries express piRNAs (Brito et al., unpublished), these data suggest that the piRNA pathway efficiently downregulates transposable and repetitive sequences in *R. prolixus* oogenesis. For all of the transposons, the RNA levels in PVS and Egg stages are generally comparable, thus pointing to a mechanism whereby transposon transcripts are expressed in the trophocytes and transported from the tropharium onto the oocyte through the trophic cords. Although our analysis does not allow us to draw conclusions on the transcriptional activity of the transposons in the follicle cells versus the germline in *R. prolixus*, the fact that transcripts corresponding to all known transposon families are detected both in PVS and in Egg stages demonstrates that none of these families is restricted to the follicle cells. It remains to be elucidated whether transposons can be transcribed in the follicle cells and their transcripts transferred to the oocyte during oogenesis or rather the transcription of these repetitive sequences occurs both in somatic and germ cells.

Chagas disease is a main threat to human health worldwide, but cures or vaccines for this illness are not yet available. The most promising strategies to reduce its diffusion rely on targeting triatomine vectors. Our study sheds light on oogenesis, early embryogenesis, and adult female fertility in *R. prolixus*, a primary vector of the Chagas disease in Central and South America. Together with substantial improvements to genome annotation in this species and novel bioinformatic resources, we provide a framework for future genetic and genomic studies and for the development of novel *R. prolixus* population control or replacement strategies.

## Methods

### *R. prolixus* handling, total RNA extraction, and library preparation

The *R. prolixus* colony was raised at 28°C and 75% relative humidity and regularly fed white rabbit blood at 3-week intervals in the laboratory of Insect Biochemistry at the Institute of Medical Biochemistry at the Federal University of Rio de Janeiro, Brazil. All animal care and experimental protocols were conducted in accordance with the guidelines of the Committee for Evaluation of Animal Use for Research (Universidade Federal do Rio de Janeiro, CAUAP-UFRJ) and the NIH Guide for the Care and Use of Laboratory Animals (ISBN 0-309-05377-3). Protocols were approved by CAUAP-UFRJ under registry #IBQM155/13. Dedicated technicians in the animal facility localized at the Instituto de Bioquímica Médica Leopoldo de Meis (UFRJ) carried out all protocols related to rabbit husbandry under strict guidelines, with supervision of veterinarians to ensure appropriate animal handling. Approximately, 100 mature chorionated eggs were dissected in ice-cold Phosphate Buffered Saline (PBS) from 5 adult females two weeks after a blood meal, and RNA-Seq libraries were produced as previously described^36^. Briefly, total RNA was extracted with TRIzol Reagent (Life Technologies) according to manufacturer instructions, treated with TURBO DNA-free kit (Ambion), and subjected to paired-end RNA sequencing (RNA-Seq) library production (Illumina). The libraries were generated and sequenced on HiSeq Illumina platforms at Lactad Facility (University of Campinas, Brazil) as previously described^36^. PVS libraries were generated previously and are available from NCBI Sequence Read Archive (SRA)^36^.

### Pre-processing and genomic mapping of the RNA-Seq reads

FastQC^61^ was adopted to check the quality of the libraries in regard to per-base sequence quality, sequence duplication level, overrepresented sequences, GC content, and presence of Illumina adapters. Raw read sequences were pre-processed by filtering out read sequences with less than 20 bp in length (--length 20); removing Ns from the read ends (--trim-n); trimming Illumina adapters (--illumina) and low-quality bases (-q 20) by Trim Galore! version 0.6.1^62^. A genome index was built with the genome assembly of *R. prolixus* RproC3^5^ and *R. prolixus* Tc1-*mariner* consensus sequences^53^ for the first-pass genomic mapping. Read alignment was performed by STAR (Spliced Transcripts Alignment to a Reference) version 2.7.2b^63^ reporting read-pairs with at most 30 multiple alignments (--outFilterMultimapNmax) and a ratio between mismatches and mapped bases less or equal to 0.1 (--outFilterMismatchNoverLmax). To rescue unmapped reads in this first-pass genomic mapping, we trimmed these sequences for poly-A/T tails (tail length > 4) and we filtered by a minimum length (>25nt) and GC-content (>20%) using prinseq-lite v0.20.4 ^64^ and re-mapped again (Supplementary Table S1).

### Transcriptome improvements in the reference annotation

Uniquely mapped read-pairs were used to improve the transcriptome landscape by genome-guided transcriptome assembly using Cufflinks pipeline version 2.2.1^39^ with the following parameters: at least 20 mapped read-pairs to support an assembled transcript (--min-frags-per-transfrag 20), novel isoform abundance ≥ 40% of the major isoform abundance (--min-isoform-fraction 0.4), reference assembly RproC3 (*Rhodnius*-*prolixus*-CDC_SCAFFOLDS_RproC3.fa) was used for bias detection (-b) and to exclude artifacts (e.g. repetitive elements) (-s); and the reference gene annotation of RproC3 (*Rhodnius*-*prolixus*-CDC_BASEFEATURES_RproC3.3.gtf) was used to guide the assembly, to compare with reference genes and to classify the assembled elements. We considered the assembled elements with a complete match of intron chain (class code “=”), potentially novel isoform (class code “j”), and unknown/intergenic transcript (class code “u”). Novel isoform abundances were calculated by Salmon v1.2.1^65^ and assembled isoforms with estimated CPM < 1 (Counts per Million) on both previtellogenic and egg stage were removed from the transcriptome assembly. Transdecoder v5.5 (https://github.com/TransDecoder) was applied to predict ORF candidates in the reference transcripts and to compare with the predicted ORFs in the improved transcriptome (Supplementary Tables S2 and S3).

### Discovery of novel transcripts and repetitive elements by *de novoassembly*

Unmapped read-pairs were used for *de novo* transcriptome assembly with the Trinity method v2.5.1^38^ with default parameters to identify novel transcripts that were not previously detected due to incomplete sequencing and assembly of the *R. prolixus* genome for each development stage, PVS and egg. *De novo* assembled transcripts were post-processed to filter out potential artifacts based on abundance and quality. First, expression levels of the *de novo* assembled transcripts were quantified by Salmon v1.2.1^65^ and isoforms with TPM < 10 were removed. Next, deduplication of redundant contig sequences was performed by CD-HIT-EST v4.6^66,67^ at a nucleotide identity of 95%. TransRate software version 1.0.3^68^ used the unmapped reads and the remaining contigs from the previous filtering steps as input to evaluate common assembly errors (chimeras, structural errors, incomplete assembly, and base errors) and to produce a diagnostic quality score for each contig, thereby removing possible artifacts. Assembled transcripts remaining from this last step were considered to be good quality transcripts (Supplementary Table S4). To identify potential protein-coding transcripts, we compared the assembled contigs against the protein databases of *R. prolixus, D. melanogaster, Caenorhabditis elegans* and *Homo sapiens* from Uniprot release 2020_01^69^ using BLASTX v2.9 (e-value < 1e-10). Uniprot protein coverage was calculated using the scripts blast_outfmt6_group_segments.pl and blast_outfmt6_group_segments.tophit_coverage.pl provided by Trinity ^38^. In addition, we performed predictions of candidate coding regions (at least 100 amino acids in length) using Transdecoder v5.5 (https://github.com/TransDecoder). ORF candidates can be classified by their completeness into complete, partial (5’ or 3’) and internal. If more than one type of ORF is detected in a contig, we assign the most complete type (Supplementary Table S5). *De novo* assembled transcriptomes of PVS and eggs were deduplicated using CD-HIT-EST (identity > 95%) to produce a final non-redundant set of transcripts. Putative protein-coding transcripts with an estimated number of reads lower than 20 were removed. To identify putative transposable elements, we performed homology searches using HMMER v3.1b2 (--cut_ga) with profile hidden Markov models (HMMs) of repetitive DNA elements from Dfam database release 3.1^54,55^. We verified the uniqueness of the putative transposon sequences by comparing (using BLASTN v2.9) with the library of repetitive elements (*Rhodnius*-*prolixu*s-CDC_REPEATS.lib) of the assembly RproC3 available in VectorBase ^5^ and consensus sequences from Tc1-*mariner*^53^ (Supplementary Table S10).

### Read coverage of the improved gene annotation and repetitive elements

Read coverage of protein-coding genes was assayed by counting the number of genes of the reference and improved gene annotation overlapped by uniquely mapped read-pairs (at least 40 mapped read-pairs) using featureCounts version 2. We set featureCounts to only consider read-pairs that have both ends aligned and to not include read-pairs that have their ends mapping to different chromosomes or mapping to the same chromosome but on different strands (-p -B -C --largestOverlap -O). To quantify the read coverage of the repetitive element classes, we included the multi-mapped read-pairs in the read assignment (--fraction -M) to the RepeatMasker and improved gene annotation of *R. prolixus*. Read-pairs assigned to more than one class of genomic elements (protein-coding gene, transposable element, simple repeat, multi-copy genes, etc.) were considered ambiguous. Reference gene annotation, RepeatMasker annotation, and genome assembly of the RproC3 are available at VectorBase^5^. To reduce the ambiguity in the read-pair assignment, we merged the overlapped repetitive elements from the same class and strand (Supplementary Table S9). Total coverage for each repetitive element class was calculated by summing the partial coverage of individual elements of the same class. We considered TE merged regions longer than 0.5 Kb^5^.

### Quantification and differential gene expression analysis

Quantification of the improved gene annotation was performed by Salmon 1.2.1^65^, which was set to produce aggregated gene-level abundance estimates (-g). The transcriptome index was built using k-mer size equal to 19 (-k 19) and the DNA sequences of the improved transcriptome. These sequences were extracted using the script gtf_genome_to_cdna_fasta.pl (https://github.com/TransDecoder). The estimated number of reads was used for the differential gene expression analysis. Genes with low expression levels were removed; we retained genes with counts scaled by total mapped reads per million (CPM) greater than 1 in at least two samples for differential gene expression analysis. We applied Trimmed Mean of M values^70^ to normalize gene counts by sequencing depth and RNA composition. DESeq2^42^ was applied to evaluate the differential gene expression during the transition of the two developmental stages using DEBrowser v1.16.1^71^. Genes were considered differentially expressed when they had an adjusted p-value ≤ 0.01 (FDR); fold-change ≤ −2.5 (most abundant in PVS) or fold-change ≥ 2.5 (most abundant in egg data sets). Differentially expressed genes with negative or positive fold-change are labeled as “down-regulated” and “up-regulated” genes, if their expression is respectively lower or higher in eggs versus PVS stages. Heatmaps of the top 25 most differentially expressed genes were generated by Heatmapper^72^ using the normalized expression (CPM) for each sample. Gene product annotation was obtained from Uniprot^69^.

### Enrichment analysis

We compared the enriched functional categories annotated for differentially expressed genes of each stage. We used Uniprot protein’s best hits (based on score and e-value) of *R. prolixus* assigned to the DNA sequences of the *R. prolixus* transcripts by BLASTX v2.9 ^73^ (-evalue 0.05) as the input for gene ontology enrichment analysis using g:Profiler (Reimand et al. 2016). All known *R. prolixus* genes were used as statistical domain scope or background. GO terms were considered enriched when Benjamini-Hochberg FDR ≤ 0.05. ClueGO ^44,45^ was used to fuse the GO parent-terms based on similar associated genes and produce functionally grouped networks (Supplementary Fig. S3).

## Acknowledgments

We are grateful to Yuri Pritykin, Julie Merkle, Olivier Devergne and Rodrigo Nunes da Fonseca for critical reading of the manuscript and to Antonio Bernardo de Carvalho, Márcia Cury El-Cheikh, Pedro Lagerblad de Oliveira, and Rafael Mesquita for invaluable help and constant technical support. We thank Graciela Venturi for assistance with confocal microscopy.

## Author contributions

AP conceived the study. VC, TB and AP performed the bioinformatic analyses. MS developed the RIO browser with inputs from AP. IB, MC and MB carried on the wet biology experiments. MC and MB supervised and oriented younger students involved in the project. VC, TB, MC, MB, HA and AP contributed to the interpretation of the data. HA and AP provided financial support. VC and AP wrote the manuscript with input and help from all the authors. The authors declare no competing interests.

## Funding Statement

This work was supported by the National Counsel of Technological and Scientific Development (CNPq) (MS and AP), the Research Support Foundation of the State of Rio de Janeiro (FAPERJ) (HA, MS and AP), INCT/Enem (HA), and a Wellcome Trust grant 207486Z17Z (AP). Master’s and Ph.D. fellowships for TB, IB, and MB were provided by CNPq. Postdoctoral fellowships for VC and MC were provided by Wellcome-Trust and Coordenação de Aperfeiçoamento de Pessoal de Nível Superior (CAPES), respectively. The funders had no role in study design, data collection and analysis, decision to publish, or preparation of the manuscript.

## Data availability

RNA-Seq datasets can be accessed at SRA NIH, accession number …. ; all other data can be found in the manuscript and supporting information files. A visual representation of the improved transcriptome that we elaborated in our work is available at http://www.genome.rio/cgi-bin/hgGateway?db=rproC3.

## Notes

### Competing Interest Statement

The authors have declared no competing interest.

